# Brain Age Gap Associations with Externalising Behaviours in Youth: A Longitudinal Investigation in the Adolescent Brain Cognitive Development (ABCD) Cohort

**DOI:** 10.64898/2026.10.22.761283

**Authors:** A. Asikin, R. Visontay, E. K. Devine, L. M. Squeglia, H. Byrne, L. Mewton

## Abstract

**Background:** Early adolescence is a period of heightened risk for the emergence of externalising psychopathology, making it an important window for identifying neurodevelopmental risk markers. Differences between predicted brain age and chronological age–termed the *brain age gap*–reflect variation from age-typical patterns of brain structure.

While brain age gap has been closely linked with internalising behaviours in adolescence, less is known about its relationship with externalising symptoms. This study examined whether brain age gap in late childhood/early adolescence (9-11 years) is associated with current externalising symptoms and whether it predicts externalising trajectories over seven- year follow-up.

**Methods:** Participants were 10,516 adolescents (aged 9-11 at baseline; up to 18 years at follow-up) from the Adolescent Brain Cognitive Development (ABCD) Study who met inclusion/exclusion criteria. Brain age gap for each participant was derived from an external validated prediction model (Drobinin et al., 2022) applied to baseline measurements of cortical volume, area, and subcortical volume. Linear mixed-effects models were used to test cross-sectional and longitudinal associations between baseline brain age gap and externalising symptoms, as assessed using the Child Behaviour Checklist. Primary models were adjusted for age, sex, pubertal status, and prior to inclusion of sociodemographic covariates (ethnoracial identity and caregiver highest education).

**Results:** A higher/more positive brain age gap was associated with greater externalising symptoms at baseline (*B*[SE]=0.067[0.033], *p*=.042), but this association was removed following sociodemographic adjustment (*B*[SE]=0.056[0.033], *p*=.088). Longitudinally, a significant brain age gap × age interaction was identified that remained evident after sociodemographic adjustment, with this relationship weakening across adolescence (*B*[SE]=- 0.007[0.003], *p*=.033).

**Conclusions:** These findings suggest that a more positive brain age gap, indicating more mature-appearing brain structure for age, in late childhood/early adolescence may be associated with externalising psychopathology, but this relationship is likely to be age- and context-dependent rather than a stable marker of persistent behavioural risk.

## Background

Externalising psychopathology, characterised by outward-directed behaviours such as aggression, rule-breaking, and impulsivity, is common during childhood and adolescence and is associated with substantial individual and societal burden (Santomauro et al., 2026).

Greater externalising symptom severity in childhood has been associated with poorer psychosocial outcomes in adulthood, including increased risk for physical aggression and risky substance use (Reef et al., 2011). The emergence of externalising symptoms during this age period coincides with substantial neurodevelopmental change, characterised by rapid nonlinear changes in cortical morphology and white matter development (Tamnes et al., 2017a, 2018). This overlap has prompted interest in whether individual differences in brain maturation may contribute to variation in the severity and persistence of externalising psychopathology during this key neurodevelopmental period.

Neuroimaging studies of externalising psychopathology in youth have identified widespread structural differences, but these findings are typically examined across individual regions or morphometric features, making their broader developmental interpretation unclear. For example, while externalising symptoms in late childhood/early adolescence have been associated with lower cortical volumes (Blok et al., 2022; Mewton et al., 2022; Zhang et al., 2022), reduced cortical thinning (Whittle et al., 2020), and variation in subcortical volume development (Bos et al., 2018; Jarvers et al., 2022), it remains difficult to determine whether these findings reflect a broader pattern of age-related brain organisation. This has prompted interest in complementary approaches that integrate information across multiple brain structural features to provide a broader index of age-related variation in brain structure.

One integrative approach to characterising such changes is brain age prediction, which uses multivariate neuroimaging features and machine learning to estimate an individual’s apparent “brain age”. The difference between this predicted age and chronological age is termed the brain age gap (‘BAG’ or ‘BrainAGE’). Lower/more negative BAG values are considered to reflect relatively less “mature-appearing” brains, whereas higher/more positive values indicate relatively more mature-appearing brains for a given chronological age (Whitmore & Beck, 2025). Although emerging work has linked more positive BAG and a more positive shift in BAG over time with internalising psychopathology, including major depressive disorder and depressive and anxiety symptoms (Drobinin et al., 2022; MacSweeney et al., 2025), its relationship with externalising psychopathology remains comparatively underexplored. Exploring this question is particularly important given the strong developmental links between externalising symptoms and harmful behaviours such as substance use, for which early risk identification may inform prevention efforts. Notably, our recent work found that a more negative BAG in late childhood was associated with subsequent alcohol initiation in adolescence, although this relationship appeared to partly reflect broader sociodemographic and prior behavioural influences (Byrne et al., 2026).

Given that childhood and adolescent externalising symptoms are associated with a 62% greater risk of later alcohol use disorder (Meque et al., 2019), it is therefore possible that BAG–alcohol associations may reflect a broader externalising liability.

Using data from the Adolescent Brain Cognitive Development (ABCD) Study (release 7.0; N = 11,860, 9-11 years at baseline), the present study investigated associations between estimated BAG and externalising psychopathology. Specifically, we examined whether deviations in brain age during late childhood predict concurrent externalising, as well as externalising into late adolescence.

## Methods & Materials

### Participants

Data from the ongoing longitudinal ABCD Study (release 7.0) were used. The ABCD Study is the largest long-term brain development and child health study to be conducted in the United States. Participants were recruited from 21 sites across the United States, primarily through schools selected based on their demographic composition, including gender, race/ethnicity, socioeconomic status, and urbanicity (Garavan et al., 2018). A central Institutional Review Board for the study protocols was granted by the University of California, San Diego, as well as from individual review boards at each study site. Before participating in the ABCD Study, written informed consent was obtained from a parent or guardian, and assent from the participants (Garavan et al., 2018). For secondary analysis of de-identified ABCD data in the current study, no additional ethical approval was required. All methods were performed in accordance with the relevant guidelines and regulations.

ABCD Release 7.0 included 11,860 participants at baseline. For BAG estimation, participants were excluded for falling outside the predefined 9-11 year age window (n = 205), unavailable baseline structural imaging data (n = 107), failing predefined MRI quality control inclusion criteria (n = 195; Hagler et al., 2019), or missing pubertal status (n = 837), leaving n = 10,516 participants eligible for harmonisation and subsequent BAG estimation. Two participants with missing family ID and two with missing baseline externalising data were excluded, leaving N = 10,512 for primary cross-sectional analyses. Secondary models included N = 10,499 due to additional missing caregiver education data. For the longitudinal models, externalising data were available from baseline through the 7-year follow-up, corresponding approximately to ages 9-18 years. All participants with BAG estimates had at least one available externalising assessment; two with missing family ID were excluded, yielding N = 10,514 in the primary longitudinal model. Secondary models included N = 10,501, reflecting an additional 13 participants with missing caregiver education data.

### Image processing

Preprocessed baseline structural MRI data (ages 9-11 years) were used (Hagler et al., 2019). Briefly, cortical parcellation and subcortical segmentation of T1-weighted images were performed using FreeSurfer v7.1.1 (https://surfer.nmr.mgh.harvard.edu/). Cortical volume and surface area measures were mapped to the Desikan-Killiany atlas, comprising 34 cortical regions per hemisphere (68 regions total; Desikan et al., 2006), alongside 53 global and subcortical volumetric measures derived from FreeSurfer *aseg* outputs.

To reduce scanner-related technical variability while preserving biologically relevant variation, imaging measures were harmonised across scanners using *ComBat* (v1.0.13; Fortin et al., 2018). Scanner device serial number was modelled as the batch variable, while chronological age, sex, and pubertal status (derived from caregiver-reported Pubertal Development Scale scores; see *Covariates*) were included as biological covariates to be preserved during harmonisation, as performed in previous work (Byrne et al., 2026).

### BAG estimation

BAG was estimated using the pretrained developmental *XGBoost* model by Drobinin et al. (2022), which was applied directly to the ABCD data. This model was selected based on its favourable performance and external validation in paediatric cohorts (Michael et al., 2026), and to maintain methodological consistency with previous work (Byrne et al., 2026). The original model was trained on 1,299 typically developing participants aged 9-19 years and achieved a mean absolute error (MAE) of approximately 1.5 years in validation samples. It has also been independently evaluated in baseline ABCD participants aged 9-11 years, with a raw MAE of 2.32 years and bias-adjusted MAE of 1.45 years (Whitmore et al., 2023).

BAG estimation followed the ABCD implementation described by Whitmore et al. (2023) and Byrne et al. (2026). Of the 189 FreeSurfer-derived structural features used by the original model, 175 were available in ABCD; the remaining 14 comprised global summary and small peripheral structure measures and were coded as missing, which is handled natively by *XGBoost*. The pretrained model was applied without retraining to preserve independence from the ABCD sample and ensure comparability with previous work. A full list of included features can be found at: https://github.com/Lifespan-and-brain-health-team/BAG-externalising/.

### Bias correction

BAG estimates were corrected for systematic age-related prediction bias using the slope and intercept parameters derived from the Drobinin et al. (2022) validation procedure, consistent with previous applications of the model in ABCD (Byrne et al., 2026; Whitmore et al., 2023). Chronological age was additionally included as a covariate in all statistical models to account for residual age-related bias, as recommended in other work (Peng et al., 2021; Whitmore & Beck, 2025).

### Externalising

Youth externalising psychopathology was obtained from the externalising subscale of the Child Behaviour Checklist caregiver-report questionnaire for youth aged 6 to 18 years (CBCL/6-18; Achenbach & Ruffle, 2000). The externalising subscale comprises aggression and rule-breaking behaviour related items, which are rated on a 3-point Likert scale (0: Not true, 1: Somewhat or sometimes true, 2: Very true or often true) based on the behaviours of the preceding 6-months. The summed value of the items provides a total Externalising score, where higher scores represent greater externalising symptoms. In the current study, raw summed scores were used rather than age- and sex-standardised T-scores to preserve developmental variation across adolescence. The externalising score was derived at baseline and over seven follow-ups (eight assessment occasions in total) acquired at yearly intervals.

### Covariates

Age, sex, and pubertal status at baseline were included as primary covariates based on prior associations with BAG estimates (Holm et al., 2023; Whitmore et al., 2023). Chronological age was collected at each timepoint and sex at birth was determined at baseline from parent report. Pubertal status was assessed using the caregiver-reported Pubertal Development Scale (PDS; Petersen et al., 1988), which indexes physical markers of pubertal maturation on a five-stage scale (1-5) approximating the Tanner stages, with higher scores indicating more advanced development. Caregiver-reported PDS scores were prioritised given evidence of greater uncertainty and missingness in youth self-report at the ABCD baseline assessment (Cheng et al., 2021).

As developmental BAG has been linked with socioeconomic and environmental factors, including socioeconomic disadvantage, neighbourhood context, and broader social determinants of health (including inequities associated with racism and discrimination), ethno-racial identity and primary caregiver education were included in secondary models as a proxy for these broader social and environmental domains (Noble et al., 2015; Rakesh et al., 2021; Rakesh & Whittle, 2021). Ethno-racial identity was categorised as Hispanic, White, Black, Asian, or Other according to the five-level ABCD Study classification derived from caregiver-reported race and ethnicity at baseline in accordance with NIH reporting standards.

Caregiver education was categorised across six levels ranging from less than high school to professional or doctoral degree. Importantly, these sociodemographic variables were not interpreted as biological traits, but rather as markers of broader structural and sociocultural influences that may shape neurodevelopmental and behavioural outcomes. For transparent reporting, a breakdown of the original caregiver education categories and the full 15-level ABCD ethno-racial identity classification is provided in *Table S1*.

### Statistical analysis

Statistical analyses were conducted in *RStudio* (R v4.5.3; RStudio v2026.1.2.418; R Core Team, 2024; RStudio Team, 2020). Associations between BAG and caregiver-reported externalising symptoms were examined using linear mixed-effects models implemented with the *lmer* function from the *lme4* package (v2.0.1; Bates et al., 2015), with statistical inference obtained using the *lmerTest* package (v3.2.1; Kuznetsova et al., 2017). Primary models adjusted for chronological age, sex, and pubertal status. Secondary models additionally included ethno-racial identity and primary caregiver education to assess the robustness of associations to broader sociodemographic influences.

Cross-sectional models examined associations between baseline BAG and baseline externalising symptoms. Longitudinal models examined whether baseline BAG was associated with externalising trajectories across adolescence by including time-varying chronological age and a BAG × age interaction. To account for the hierarchical structure of the data, all models included random intercepts for family nested within study site, while longitudinal models additionally included a participant-level random intercept to account for repeated observations.

### Sensitivity analysis

To assess whether baseline externalising symptoms were associated with follow-up data availability, a logistic regression tested whether baseline externalising symptoms predicted availability of wave-7 externalising data, adjusting for baseline age, sex, and pubertal status.

## Results

### Participant characteristics

Participant characteristics are summarised in *Table 1*. The final BAG sample included 10,516 participants. Flowchart of participant-level exclusions are shown in Figure 1.

**Figure 1.**
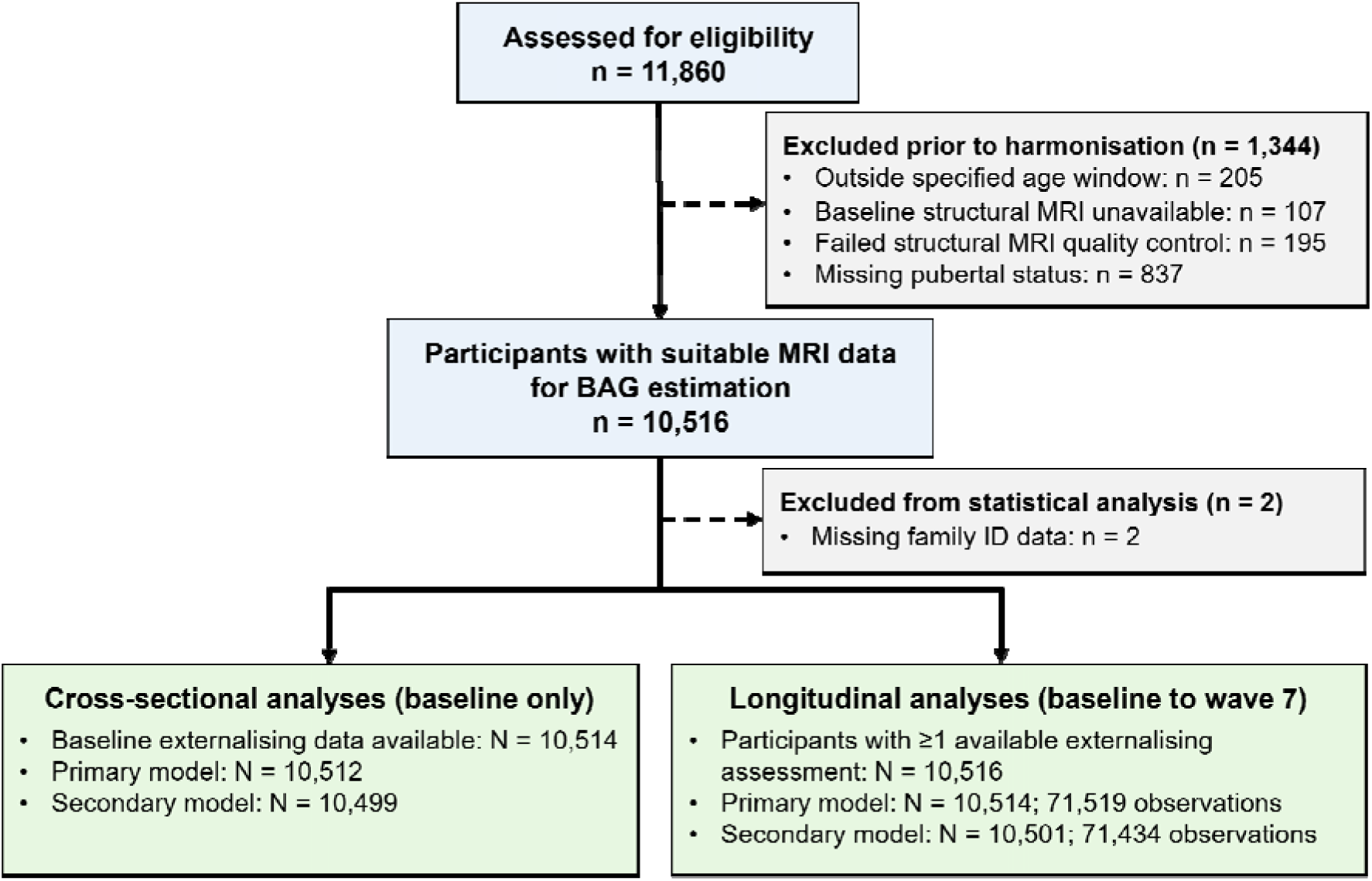
Flowchart for participant inclusion/exclusion.

**Figure 2.**
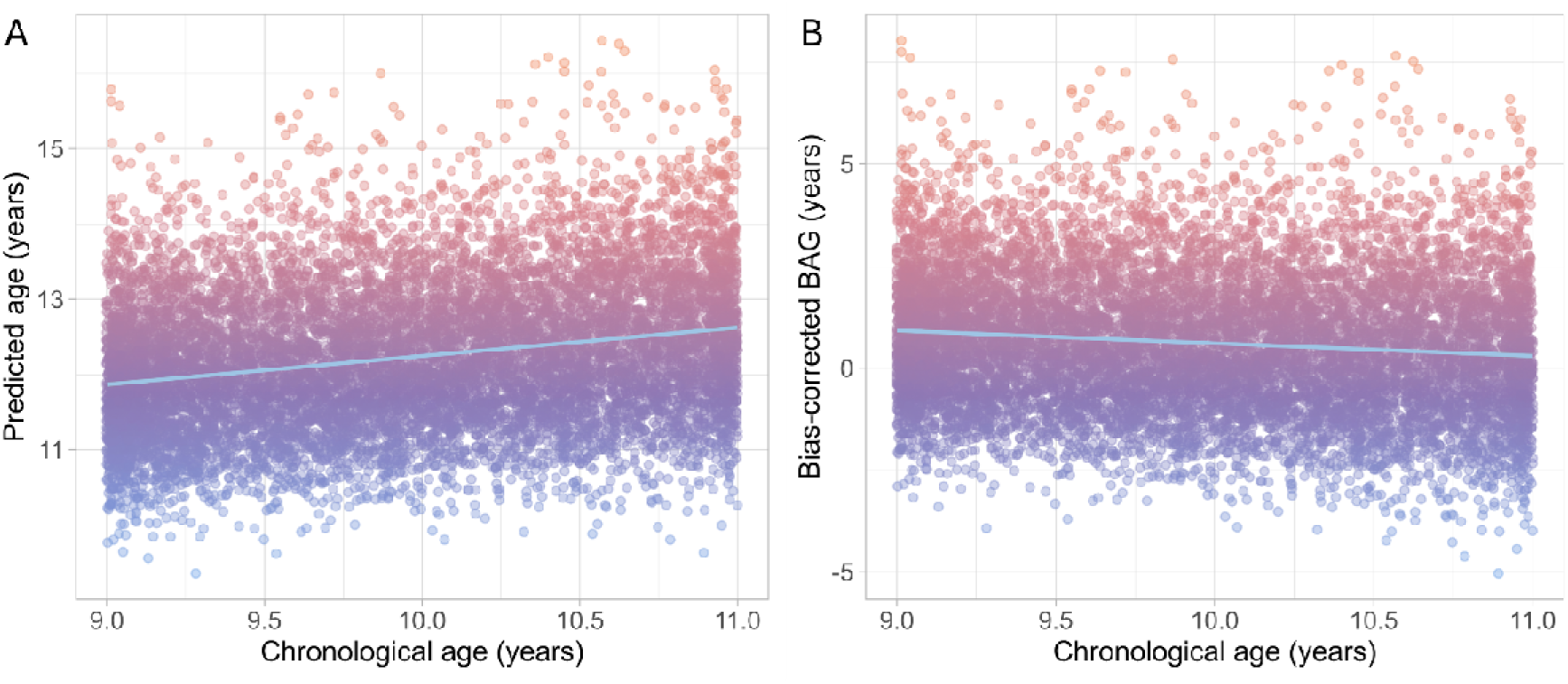
Scatterplots showing: (A) chronological age compared to raw predicted age prior to age bias correction; (B) chronological age compared to bias-corrected BAG within the full study sample. Bias corrected values were used for subsequent analyses. Warmer colours indicate a higher/more positive BAG, while cooler colours indicate lower/more negative BAG.

**Table 1.** Baseline characteristics of participants included in the final brain age gap (BAG)

| Characteristic | N = 10,516 |
| --- | --- |
| Bias-corrected Brain Age Gap (BAG)<sup>1,3</sup> | 0.62 (1.71 [-5.03, 8.03]) |
| Age<sup>1</sup> | 9.95 (0.62 [9.0-11.0]) |
| Sex<sup>2</sup> |  |
| Female | 5,235 (50%) |
| Pubertal Status<sup>2</sup> |  |
| Pre Pubertal | 5,584 (53%) |
| Early Pubertal | 2,277 (22%) |
| Mid Pubertal | 2,484 (24%) |
| Late Pubertal | 161 (1.5%) |
| Post Pubertal | 10 (<0.1%) |
| Ethnoracial Identity<sup>2</sup> |  |
| Hispanic | 2,127 (20%) |
| White | 5,714 (54%) |
| Black | 1,415 (13%) |
| Asian | 203 (1.9%) |
| Other | 1,057 (10%) |
| Highest Caregiver Education<sup>2</sup> |  |
| Professional or Doctoral Degree | 659 (6.3%) |
| Master's Degree | 2,101 (20%) |
| Bachelor's Degree | 3,064 (29%) |
| Some college or Associate's Degree | 3,048 (29%) |
| High School Graduate or GED | 1,015 (9.7%) |
| Less than High School | 616 (5.9%) |
| Missing/Unknown | 13 (0.1%) |

<sup>1</sup> Mean (SD [range]); <sup>2</sup> n (%); <sup>3</sup> Brain Age Gap (BAG) refers to difference between predicted age and chronological age. Values less than zero are considered to reflect relatively less “mature-appearing” brains, whereas values above zero indicate relatively more mature-appearing brains for a given chronological age.

**Table 2.** Longitudinal age and externalising symptom characteristics across assessment waves.

| Wave | Age<sup>1</sup> | Externalising<sup>1</sup> | *n* available | Missing/Unknown |
| --- | --- | --- | --- | --- |
| Baseline | 9.95 (0.62) | 4.41 (5.82) | 10514 | <10 (<0.1%) |
| 1-year follow-up | 10.96 (0.64) | 4.13 (5.57) | 9981 | 535 (5.1%) |
| 2-year follow-up | 12.07 (0.66) | 3.89 (5.56) | 9727 | 789 (7.5%) |
| 3-year follow-up | 12.96 (0.64) | 3.90 (5.51) | 9145 | 1371 (13.0%) |
| 4-year follow-up | 14.18 (0.70) | 3.62 (5.47) | 8537 | 1979 (18.8%) |
| 5-year follow-up | 15.08 (0.66) | 3.77 (5.76) | 7854 | 2662 (25.3%) |
| 6-year follow-up | 16.06 (0.68) | 3.24 (5.27) | 8358 | 2158 (20.5%) |
| 7-year follow-up | 16.94 (0.66) | 3.26 (5.23) | 7415 | 3101 (29.5%) |

<sup>1</sup> Mean (SD).

### Model performance

Consistent with previous applications of the model in ABCD (Whitmore et al., 2023; Byrne et al., 2026), the pretrained model showed systematic age-related prediction bias. Prior to correction, predicted age exceeded chronological age by an average of 2.27 years (SD = 1.01), with an MAE of 2.28 years. Raw predicted age correlated positively with chronological age (*r* = .24, 95% CI [.22, .26], *p* < .001), whereas raw BAG was negatively correlated with chronological age (*r* = -.38, 95% CI [-.39, -.36], *p* < .001). Following linear bias correction, MAE improved to 1.42 years and mean BAG was +0.62 years (SD = 1.71). Bias-corrected BAG estimates were used in all subsequent analyses.

## BAG Associations with Externalising Psychopathology

### Cross-sectional (Baseline only)

At baseline (ages 9–11 years), a higher brain age gap was associated with greater caregiver-reported CBCL externalising symptoms after adjusting for age, sex, and pubertal status (*B* [SE] = 0.068 [0.033], *p* = .041; *Figure 3*). Following additional adjustment for ethno-racial identity and primary caregiver education, this association was no longer statistically significant (*B* [SE] = 0.057 [0.033], *p* = .083). Full model results are reported in *Table S2*.

**Figure 3.**
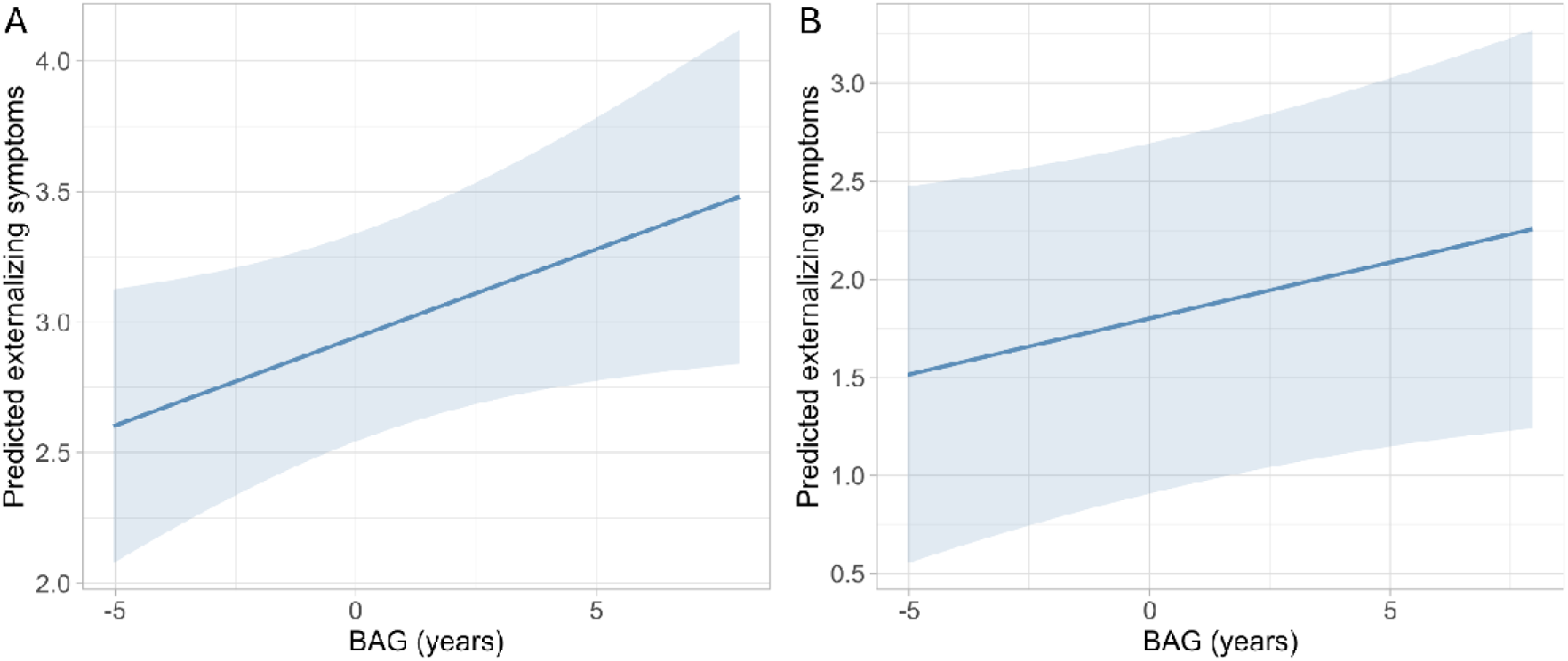
Predicted caregiver-reported CBCL externalising symptoms according to baseline BAG. (A) Reduced model adjusted for age, sex, and pubertal status. (B) Full model additionally adjusted for ethno-racial identity and highest caregiver education. Lines represent model-predicted externalising symptoms across BAG; shaded areas indicate 95% confidence intervals.

### Longitudinal (Baseline to Wave 7)

In the longitudinal models, a significant BAG × age interaction indicated that the positive association between baseline BAG and externalising symptoms weakened with increasing age (*B* [SE] = -0.007 [0.003], *p* = .036). This interaction remained significant following additional adjustment for ethno-racial identity and primary caregiver education (*B* [SE] = - 0.007 [0.003], *p* = .039; *Figure 4*). Full model results are reported in *Table S3*.

**Figure 4.**
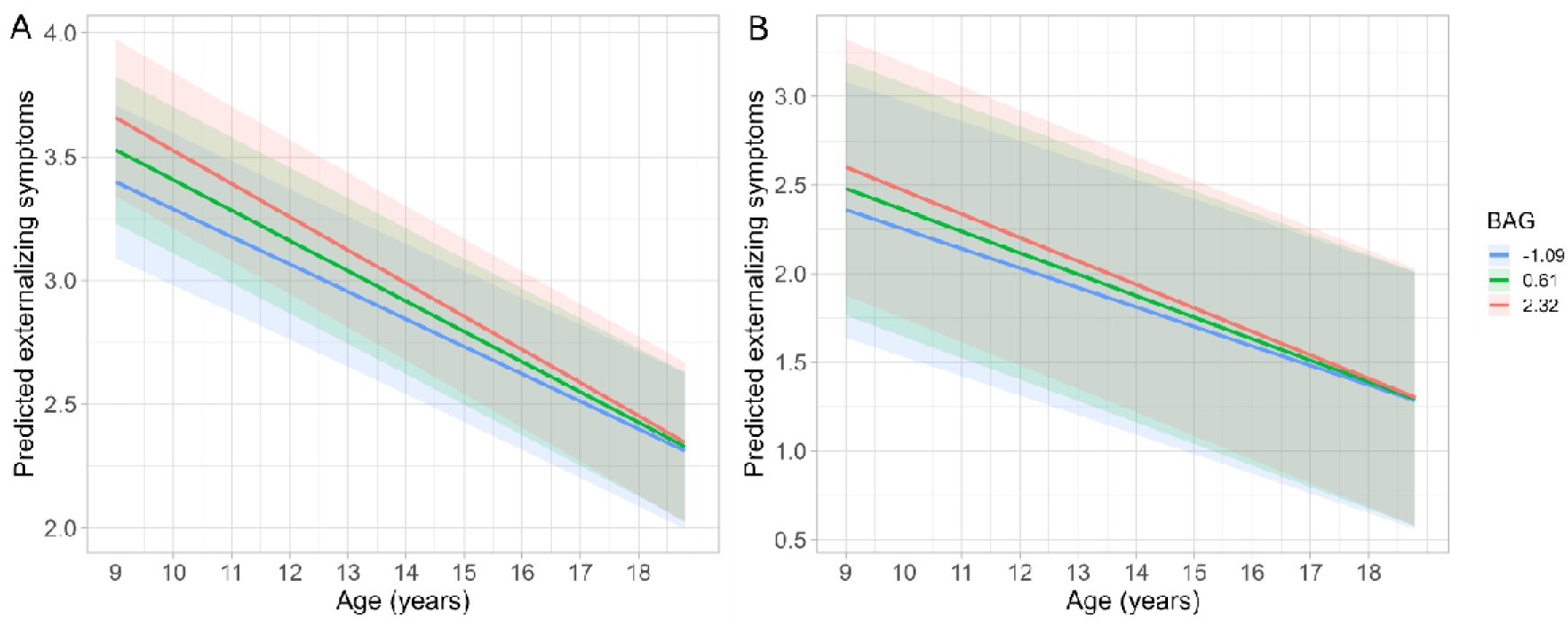
Predicted trajectories of caregiver-reported CBCL externalising symptoms across chronological age according to BAG. (A) Reduced model adjusted for sex and pubertal status. (B) Full model additionally adjusted for ethno-racial identity and caregiver education. Both models included chronological age and the BAG × age interaction. Lines represent low (-1 SD), mean, and high (+1 SD) baseline BAG; shaded areas indicate 95% confidence intervals.

### Sensitivity analysis

Baseline externalising symptoms were modestly higher among participants without wave-7 data (mean [SD] = 5.07 [6.33]) than among those with wave-7 data (mean [SD] = 4.13 [5.56]). Higher baseline externalising was also associated with lower odds of wave-7 data availability after adjustment for baseline age, sex, and pubertal status (OR = 0.975, 95% CI [0.968, 0.982], *p* < .001).

## Discussion

Using data from the ABCD Study, we examined whether BAG measured in late childhood– early adolescence (ages 9-11) was associated with concurrent externalising symptoms and their trajectories through late adolescence. At baseline, a more positive BAG, reflecting more mature-appearing brain structure for chronological age, was associated with greater externalising symptoms, although this association was no longer evident after sociodemographic adjustment. In contrast, the association between baseline BAG and externalising symptom trajectories remained evident after adjustment but weakened across adolescence, with predicted symptom trajectories converging by late adolescence. Together, these findings suggest that BAG in late childhood-early adolescence may capture age- and context-dependent variation in externalising psychopathology rather than a stable marker of persistent risk. Future studies should replicate both the cross-sectional and longitudinal findings in independent cohorts, ideally using repeated brain-age estimates and alternative estimation models, to assess their generalisability beyond the symptom decline and selective attrition observed in ABCD.

The association between externalising symptoms and a more mature-appearing structural profile is broadly consistent with previous work in ABCD. Greater externalising symptoms have been linked to smaller cortical volumes and lower surface area at baseline (Mewton et al., 2022; Zhang et al., 2022), features that typically decline across adolescence (Tamnes et al., 2017b). By integrating these feature-specific associations within a single age-related metric, the present BAG findings extend this work into a broader whole-brain context.

However, considerable caution is warranted when interpreting brain-age estimates in adolescence, as BAG represents a summary deviation from age-typical brain structure rather than a direct measure of maturational change (Whitmore & Beck, 2025), particularly when estimated at a single timepoint. Other features also complicate this interpretation; externalising associations with subcortical volumes appear to vary across development (Bos et al., 2018; Jarvers et al., 2022), and cortical thickness, which was not included in the present brain-age model, has been linked to patterns consistent with attenuated maturation (Whittle et al., 2020). This divergence highlights the difficulty of summarising heterogeneous structural features in a single brain-age metric, especially when feature-level contributions are not readily interpretable. Future work combining longitudinal BAG with multimodal, connectomic, and network-based approaches may clarify which aspects of brain organisation underlie these associations.

A related challenge is that little is known about how environmental and contextual factors shape brain-age variation in adolescence. This is particularly relevant here, as the attenuation of the concurrent association after sociodemographic adjustment suggests that contextual influences may shape this relationship. This is consistent with our previous work in ABCD, in which an initial association between baseline BAG and alcohol initiation by mid-adolescence was largely explained by similar contextual factors (Byrne et al., 2026).

However, these characteristics – specifically, ethno-racial identity and highest caregiver education – are only coarse proxies for broader contextual exposures, and their contribution is therefore likely to be underestimated. More broadly, cross-sectional and longitudinal studies have linked BAG estimates to socioeconomic disadvantage, neighbourhood context, and wider social determinants of health (Cohen et al., 2024; Rakesh et al., 2021; Rakesh & Whittle, 2021; Yang et al., 2026), although the direction of these associations appears to be context- and age-dependent. These broader contextual influences have also been implicated in externalising psychopathology (Kim et al., 2022; Peverill et al., 2021), suggesting that shared contextual factors may underlie variation in both BAG and symptom burden and partly account for the cross-sectional association observed here. Together, these findings highlight the need to better understand how environmental and contextual influences contribute to variation in BAG and its associations with psychopathology, to better characterise the processes captured by developmental brain-age measures.

In contrast to the cross-sectional findings, the longitudinal BAG-externalising relationship remained robust after sociodemographic adjustment, but weakened with increasing age.

Specifically, youth with higher baseline BAG showed greater externalising symptoms earlier in adolescence, but these differences diminished over time, with predicted trajectories for higher- and lower-BAG youth converging by approximately ages 17-18. This pattern indicates that BAG-related differences in externalising symptoms are not stable across adolescence and may be most prominent around the developmental period in which BAG was assessed. However, because brain age was calculated only at baseline, it remains unclear whether this convergence reflects subsequent changes in brain development or other developmental, environmental, or psychosocial processes that become increasingly relevant across adolescence (Rakesh et al., 2021; Yang et al., 2026). While these findings suggest that the longitudinal relevance of BAG measured in late childhood/early adolescence may be limited, replication using longitudinal brain-age estimates, alternative brain-age models, and samples spanning different developmental periods is warranted.

The apparent convergence in externalising symptoms across adolescence should also be interpreted in the context of broader symptom trajectories within ABCD. Recent ABCD work has shown an overall decline in externalising symptoms across adolescence, while also identifying substantial heterogeneity in individual trajectories, including multiple decreasing subgroups and a smaller increasing subgroup (Brieant et al., 2026). Such heterogeneity is consistent with broader developmental research showing that some externalising symptoms decline over time, while others, including conduct problems, aggression, and rule-breaking, may emerge or increase during early-to-mid adolescence (Atherton et al., 2018; Bos et al., 2018). Accordingly, the stronger decline observed among youth with higher baseline BAG may partly reflect variation within a broader downward trend in ABCD rather than a distinct protective effect of higher BAG. Our sensitivity analysis also indicated that participants without wave-7 data had somewhat higher baseline externalising symptoms, suggesting some selective loss of participants with greater initial symptom burden at later follow-up. Together, these findings suggest that the observed convergence should be interpreted within broader developmental and sample-specific patterns of externalising symptoms, rather than as a BAG-specific effect in isolation.

### Limitations

There are multiple strengths of this study, including the use of a large, well-characterised longitudinal cohort, an externally trained and validated brain-age model, and repeated assessment of externalising psychopathology across adolescence. However, several limitations warrant consideration. First, brain-age approaches reduce complex, regionally heterogeneous patterns of neurodevelopment to a single summary index, which may limit neurobiological interpretability with respect to externalising outcomes (Whitmore & Beck, 2025). Longitudinal estimation of brain age may provide additional information by distinguishing stable individual differences in apparent brain maturity from changes in BAG across development. Further, the present model was based on structural T1-weighted MRI and therefore does not capture complementary aspects of white matter microstructure or functional organisation; future work incorporating diffusion imaging and other multimodal features may provide a more comprehensive characterisation of neurodevelopmental variation (MacSweeney et al., 2025). Finally, MRI quality-control exclusions may introduce selection bias if participants with greater externalising symptoms are disproportionately excluded because of motion. Prior ABCD work has reported clinical differences between participants included and excluded following structural imaging quality-control procedures (Mewton et al., 2022), although associations between head motion and externalising symptoms have not been consistently observed (Shi et al., 2025). Future studies should therefore examine whether psychopathology differs systematically between participants with and without usable imaging data.

There are also limitations relating to the application of the Drobinin et al. (2022) model in the present context, several of which have been discussed previously in our application of this model to ABCD data (Byrne et al., 2026). Briefly, 14 features from the original model were unavailable in ABCD, and the bias-correction parameters were derived from an external validation sample spanning a broader developmental age range (9-19 years) than the ABCD baseline cohort (9-11 years). Recent work further highlights the importance of this latter issue, demonstrating that the age distribution of BAG training samples can systematically influence prediction bias and downstream brain-behaviour associations (Konowski et al., 2026).

Although bias correction was applied and chronological age was included in all statistical models, these procedures may not fully account for biases arising from differences between the training and target age distributions. Together with the reduced feature set, this may have contributed to the tendency toward chronological age overestimation observed here.

Accordingly, BAG estimates are best interpreted as relative deviations from age-normative brain structure rather than absolute measures of biological maturity, and replication using alternative developmental brain-age models will be important.

## Conclusion

In conclusion, although higher BAG at ages 9-11 was associated with externalising symptoms both concurrently and over time, the concurrent association was no longer evident after sociodemographic adjustment, while the longitudinal association remained but weakened across adolescence. These findings suggest that BAG measured during this developmental period may capture age-specific and context-sensitive variation in externalising psychopathology rather than a stable marker of persistent risk. Future work should test these associations using longitudinal and alternative brain-age models across different neurodevelopmental periods, while examining how these relationships are embedded within broader environmental and contextual influences.

## Data availability statement

The data that support the findings of this study are available from the Adolescent Brain Cognitive Development™ (ABCD) Study through the NIH Brain Development Cohorts Data Sharing Platform. Access is restricted and subject to the relevant application, data-use, and security requirements.

## Acknowledgements

Data used in the preparation of this article were obtained from the Adolescent Brain Cognitive Development™ (ABCD) Study, held in the NIH Brain Development Cohorts Data Sharing Platform. The ABCD Study® is a multisite, longitudinal study designed to recruit more than 10,000 children aged 9–10 years and follow them over 10 years into early adulthood. The ABCD Study® is supported by the National Institutes of Health and additional federal partners under award numbers U01DA041048, U01DA050989, U01DA051016, U01DA041022, U01DA051018, U01DA051037, U01DA050987, U01DA041174, U01DA041106, U01DA041117, U01DA041028, U01DA041134, U01DA050988, U01DA051039, U01DA041156, U01DA041025, U01DA041120, U01DA051038, U01DA041148, U01DA041093, U01DA041089, U24DA041123, and U24DA041147. A full list of supporters is available at https://abcdstudy.org/federal-partners.html. ABCD Consortium investigators designed and implemented the study and/or provided data but did not necessarily participate in the analysis or writing of this report. This manuscript reflects the views of the authors and may not reflect the opinions or views of the NIH or ABCD Consortium investigators.

## Ethics approval statement

The current study involved secondary analysis of de-identified data from the ABCD Study. Ethical approval for the ABCD Study was obtained from the relevant institutional review boards at participating sites, and no additional ethical approval was required for the present secondary analysis.

## Patient consent statement

Prior to participation in the ABCD Study, written informed consent was obtained from a parent or legal guardian, and assent was obtained from participating children.

## Permission to reproduce material from other sources

Not applicable. No material requiring permission from third-party sources was reproduced in this manuscript.

## Key points and relevance

- Externalising symptoms in adolescence have been associated with structural brain differences, but their broader developmental significance remains unclear.
- Using data from 10,516 participants in the ABCD Study, we examined whether brain age gap (BAG) at ages 9-11 was associated with concurrent externalising symptoms and their trajectories across adolescence.
- Higher BAG was associated with greater externalising symptoms, although the concurrent association was no longer evident after sociodemographic adjustment and longitudinal differences weakened with age.
- These findings suggest that developmental BAG may capture age-specific and context-sensitive variation in externalising psychopathology rather than a stable marker of persistent symptoms.
- Further research should examine longitudinal brain-age estimates and environmental influences to clarify how developmental brain-age measures should be interpreted.

## CRediT authorship contribution statement

**Angela Asikin:** Conceptualization, Data curation, Formal analysis, Investigation, Methodology, Visualization, Writing – original draft, Writing – review & editing. **Rachel Visontay:** Conceptualization, Investigation, Methodology, Project administration, Writing – review & editing. **Emma K Devine:** Conceptualization, Investigation, Methodology, Project administration, Writing – review & editing. **Lindsay M Squeglia:** Conceptualization, Funding acquisition, Methodology, Supervision, Writing – review & editing. **Hollie Byrne:** Conceptualization, Data curation, Formal analysis, Investigation, Methodology, Supervision, Visualization, Writing – original draft, Writing – review & editing. **Louise Mewton:** Conceptualization, Formal analysis, Investigation, Funding acquisition, Methodology, Supervision, Writing – review & editing.

