## Supporting information for "Brain Age Gap Associations with Externalising Behaviours in Youth: A Longitudinal Investigation in the Adolescent Brain Cognitive Development (ABCD) Cohort"

*Table S1*. Detailed distribution of caregiver education and ethno-racial identity in the BAG sample (N = 10,516).

| **Characteristic** | **Category** | **n (%)** |
| --- | --- | --- |
| **Highest caregiver education** | Never attended/Kindergarten only | 0 (0.0%) |
|  | 1st grade | <10 |
|  | 2nd grade | <10 |
|  | 3rd grade | <10 |
|  | 4th grade | <10 |
|  | 5th grade | <10 |
|  | 6th grade | 57 (0.5%) |
|  | 7th grade | 17 (0.2%) |
|  | 8th grade | 46 (0.4%) |
|  | 9th grade | 105 (1.0%) |
|  | 10th grade | 85 (0.8%) |
|  | 11th grade | 152 (1.4%) |
|  | 12th grade, no diploma | 138 (1.3%) |
|  | High school graduate | 804 (7.6%) |
|  | GED or equivalent | 211 (2.0%) |
|  | Some college, no degree | 1702 (16.2%) |
|  | Associate degree: Occupational, Technical, or Vocational | 754 (7.2%) |
|  | Associate degree: Academic Program | 592 (5.6%) |
|  | Bachelor's degree (ex. BA, AB, BS, BBS) | 3064 (29.1%) |
|  | Master's degree (ex. MA, MS, MEng, MEd, MBA) | 2101 (20.0%) |
|  | Professional School degree (ex. MD, DDS, DVN, JD) | 308 (2.9%) |
|  | Doctoral degree (ex. PhD, EdD) | 351 (3.3%) |
|  | Missing | 13 (0.1%) |
| **Ethno-racial identity** | None | 652 (6.2%) |
|  | White/Caucasian | 5825 (55.4%) |
|  | Western European | 284 (2.7%) |
|  | Eastern European | 51 (0.5%) |
|  | Hispanic/Latino | 1485 (14.1%) |
|  | Black/African American | 1335 (12.7%) |
|  | Afro-Caribbean/Indo-Caribbean/West Indian | 26 (0.2%) |
|  | East Asian | 199 (1.9%) |
|  | South Asian | 52 (0.5%) |
|  | Southeast Asian | 110 (1.0%) |
|  | American Indian/Alaska Native | 72 (0.7%) |
|  | Middle Eastern/North African | 26 (0.2%) |
|  | Native Hawaiian or Pacific Islander | 20 (0.2%) |
|  | Multiracial/Multiethnic | 311 (3.0%) |
|  | Other Ethnicity | 48 (0.5%) |
|  | Missing | 20 (0.2%) |
| Percentages calculated using the full BAG sample (N = 10,516). Cell counts below 10 and their corresponding percentages are suppressed. Caregiver education reflects the highest grade or degree reported by the primary caregiver. Ethno-racial identity is reported using the full ABCD 15-level classification. | | |

Cross-sectional model results

*Table S2.* Results from cross-sectional models examining associations between baseline BAG and externalising.

|  | Primary model (N = 10,512) | | | Secondary model (N = 10,499) | | |
| --- | --- | --- | --- | --- | --- | --- |
| Predictor | B (SE) | 95% CI | p | B (SE) | 95% CI | p |
| Intercept | 5.745 (0.923) | [3.937, 7.554] | <.001 | 4.166 (1.014) | [2.179, 6.153] | <.001 |
| Brain age gap | 0.068 (0.033) | [0.003, 0.132] | 0.041 | 0.057 (0.033) | [-0.008, 0.122] | 0.083 |
| Sex: Male | 1.841 (0.133) | [1.581, 2.101] | <.001 | 1.712 (0.132) | [1.452, 1.971] | <.001 |
| Age | -0.282 (0.093) | [-0.463, -0.100] | 0.002 | -0.237 (0.092) | [-0.418, -0.057] | 0.010 |
| Pubertal status: 2 | 0.698 (0.146) | [0.411, 0.984] | <.001 | 0.593 (0.146) | [0.308, 0.879] | <.001 |
| Pubertal status: 3 | 1.162 (0.167) | [0.835, 1.489] | <.001 | 0.830 (0.171) | [0.494, 1.165] | <.001 |
| Pubertal status: 4 | 1.604 (0.465) | [0.692, 2.516] | <.001 | 1.120 (0.469) | [0.201, 2.038] | 0.017 |
| Pubertal status: 5 | 0.304 (1.760) | [-3.145, 3.753] | 0.863 | -0.454 (1.750) | [-3.885, 2.978] | 0.796 |
| Ethno-racial identity: Black |  |  |  | 1.292 (0.456) | [0.399, 2.186] | 0.005 |
| Ethno-racial identity: Hispanic |  |  |  | 0.754 (0.441) | [-0.110, 1.618] | 0.087 |
| Ethno-racial identity: Other |  |  |  | 1.581 (0.452) | [0.695, 2.467] | <.001 |
| Ethno-racial identity: White |  |  |  | 1.038 (0.422) | [0.210, 1.866] | 0.014 |
| Primary caregiver education: High school graduate or GED |  |  |  | 0.730 (0.225) | [0.289, 1.171] | 0.001 |
| Primary caregiver education: Less than high school |  |  |  | 1.493 (0.279) | [0.946, 2.040] | <.001 |
| Primary caregiver education: Master's degree |  |  |  | -0.689 (0.169) | [-1.022, -0.357] | <.001 |
| Primary caregiver education: Professional or Doctoral degree |  |  |  | -0.508 (0.254) | [-1.007, -0.010] | 0.046 |
| Primary caregiver education: Some college or Associate's degree |  |  |  | 1.037 (0.158) | [0.728, 1.345] | <.001 |

Longitudinal model results

*Table S3.* Results from longitudinal models examining associations between baseline BAG and externalising.

|  | Primary model (N = 10,514,  71,519 observations) | | | Secondary model (N = 10,501;  71,434 observations) | | |
| --- | --- | --- | --- | --- | --- | --- |
| Predictor | B (SE) | 95% CI | *p* | B (SE) | 95% CI | *p* |
| Intercept | 4.545 (0.168) | [4.210, 4.880] | <.001 | 3.492 (0.372) | [2.762, 4.222] | <.001 |
| Brain age gap | 0.137 (0.050) | [0.039, 0.235] | 0.006 | 0.130 (0.050) | [0.032, 0.228] | 0.009 |
| Sex: Male | 1.249 (0.106) | [1.041, 1.457] | <.001 | 1.170 (0.106) | [0.962, 1.378] | <.001 |
| Age | -0.118 (0.006) | [-0.130, -0.107] | <.001 | -0.117 (0.006) | [-0.128, -0.106] | <.001 |
| Pubertal status: 2 | 0.365 (0.117) | [0.136, 0.594] | 0.002 | 0.313 (0.117) | [0.084, 0.542] | 0.007 |
| Pubertal status: 3 | 0.861 (0.132) | [0.602, 1.120] | <.001 | 0.671 (0.135) | [0.405, 0.936] | <.001 |
| Pubertal status: 4 | 0.638 (0.378) | [-0.103, 1.379] | 0.091 | 0.390 (0.381) | [-0.357, 1.136] | 0.306 |
| Pubertal status: 5 | -0.294 (1.420) | [-3.077, 2.489] | 0.836 | -0.747 (1.415) | [-3.521, 2.027] | 0.598 |
| BAG × age | -0.007 (0.003) | [-0.013, -0.000] | 0.036 | -0.007 (0.003) | [-0.013, -0.000] | 0.039 |
| Ethno-racial identity: Black |  |  |  | 0.966 (0.369) | [0.242, 1.690] | 0.009 |
| Ethno-racial identity: Hispanic |  |  |  | 0.703 (0.357) | [0.003, 1.403] | 0.049 |
| Ethno-racial identity: Other |  |  |  | 1.503 (0.366) | [0.785, 2.221] | <.001 |
| Ethno-racial identity: White |  |  |  | 0.954 (0.341) | [0.285, 1.624] | 0.005 |
| Primary caregiver education: High school graduate or GED |  |  |  | 0.458 (0.186) | [0.093, 0.822] | 0.014 |
| Primary caregiver education: Less than high school |  |  |  | 0.852 (0.232) | [0.398, 1.307] | <.001 |
| Primary caregiver education: Master's degree |  |  |  | -0.484 (0.139) | [-0.756, -0.212] | <.001 |
| Primary caregiver education: Professional or Doctoral degree |  |  |  | -0.296 (0.206) | [-0.700, 0.108] | 0.151 |
| Primary caregiver education: Some college or Associate's degree |  |  |  | 0.834 (0.129) | [0.580, 1.087] | <.001 |
